# Nuclear translocation of spike mRNA and protein is a novel pathogenic feature of SARS-CoV-2

**DOI:** 10.1101/2022.09.27.509633

**Authors:** Sarah Sattar, Juraj Kabat, Kailey Jerome, Friederike Feldmann, Kristina Bailey, Masfique Mehedi

## Abstract

Severe acute respiratory syndrome coronavirus 2 (SARS-CoV-2) causes severe pathophysiology in vulnerable older populations and appears to be highly pathogenic and more transmissible than SARS-CoV or MERS-CoV [1, 2]. The spike (S) protein appears to be a major pathogenic factor that contributes to the unique pathogenesis of SARS-CoV-2. Although the S protein is a surface transmembrane type 1 glycoprotein, it has been predicted to be translocated into the nucleus due to the novel nuclear localization signal (NLS) “PRRARSV”, which is absent from the S protein of other coronaviruses. Indeed, S proteins translocate into the nucleus in SARS-CoV-2-infected cells. To our surprise, S mRNAs also translocate into the nucleus. S mRNA colocalizes with S protein, aiding the nuclear translocation of S mRNA. While nuclear translocation of nucleoprotein (N) has been shown in many coronaviruses, the nuclear translocation of both S mRNA and S protein reveals a novel pathogenic feature of SARS-CoV-2.

**Author summary:** One of the novel sequence insertions resides at the S1/S2 boundary of Spike (S) protein and constitutes a functional nuclear localization signal (NLS) motif “PRRARSV”, which may supersede the importance of previously proposed polybasic furin cleavage site “RRAR”. Indeed, S protein’s NLS-driven nuclear translocation and its possible role in S mRNA’s nuclear translocation reveal a novel pathogenic feature of SARS-CoV-2.

## Introduction

The recently emerged severe acute respiratory syndrome coronavirus 2 (SARS-CoV-2), along with SARS-CoV and Middle East respiratory syndrome coronavirus (MERS-CoV), belong to the *Coronaviridae* virus family. The current ongoing outbreak has shown that SARS-CoV-2 is highly pathogenic and more transmissible than SARS-CoV or MERS-CoV [1]. These coronaviruses contain a positive-strand RNA genome with a few unique features: two-thirds of the viral RNA is translated into a large polyprotein, and the remainder of the viral genome is transcribed by a discontinuous transcription process into a nested set of subgenomic mRNAs [3-5]. The different subgenomic RNAs encode four conserved structural proteins (spike, S; envelope, E; membrane, M, and nucleocapsid, N) and several accessory proteins [6, 7]. The S protein of both SARS-CoV and SARS-CoV-2 interacts with the host cell receptor angiotensin converting enzyme 2 (ACE2) and triggers fusion between the viral envelope and host cell membrane to facilitate successful viral entry [8, 9]. However, the S protein of MERS-CoV binds to dipentidyl peptidase (DPP4) to facilitate entry into cells [10]. Importantly, the SARS-CoV-2 S protein is a significant pathogenic factor because of its broad tropism for mammalian ACE2 [11]. While the S protein is an attractive target for therapeutic development [12], the lack of comprehensive information on S protein expression and subcellular translocation hinders the identification of an effective S protein-targeting therapeutic to combat SARS-CoV-2 infection.

The genome sequence is generally the blueprint for detecting biological function [13]. Thus, the S protein’s function is encoded in the S gene sequence. Identifying novel features in the S gene sequence, its expression and subcellular localization may shed light on the unique pathogenesis of SARS-CoV-2 compared to other pathogenic beta-coronaviruses, particularly SARS-CoV and MERS-CoV. A recent study showed several SARS-CoV-2 genomic features, including novel sequence insertions and enhanced N protein nuclear localization signals (NLSs) that are thought to be responsible for the unique pathogenesis of this coronavirus [14]. There are three types of NLSs: pat4, pat7, and bipartite. The pat4 signal is a chain of 4 basic amino acids consisting of lysine or arginine or three basic amino acids, with the last amino acid being either histidine or proline. The pat7 signal begins with proline and is followed by six amino acids, which contains a four-residue sequence in which three of the four residues are basic. The bipartite signal consists of two basic amino acids with a 10-residue spacer and a five amino acid sequence in which at least three of the five amino acids are basic [15-17]. The subcellular localization of some SARS-CoV-2 proteins has been studied in vitro [18], but a comprehensive understanding of the subcellular localization of the S protein is missing.

Here, we first report the nuclear translocation of S protein and mRNA in SARS-CoV-2-infected cells. The translocation of the SARS-CoV-2 S mRNA appeared to be assisted by the S protein, which contains an NLS motif that is unique among human pathogenic beta-coronaviruses.

## Results

### The novel NLS motif “PRRARSV” is in the S protein of SARS-CoV-2 but not SARS-CoV or MERS-CoV

Several groups have reported novel nucleotide insertions in the S gene of SARS-CoV-2, as indicated by a multiple sequence alignment for the S protein sequences of different coronaviruses, such as a polybasic site “PRRA” produced by a 12-nucleotide acquisition at the S1-S2 boundary through multiple host-species adaptations [19, 20]. However, S protein sequence alignments between SARS-CoV-2 and SARS-CoV showed the possibility of the insertions “NSPR” [21] and “SPRR” [22] at the S1-S2 boundary. It has previously been reported that the sequence insertion at the S1-S2 boundary constitutes a furin cleavage site [23, 24]. A comprehensive understanding of the consequence of the sequence insertion at the S1-S2 boundary is still missing, possibly because research focused on understanding the differences in the pathogenicity of the different SARS-CoV-2 variants and subvariants, which emerged rapidly. To determine whether the earlier SARS-CoV-2 isolate (USA/WA-CDC-WA1/2020 isolate, GenBank accession no. MN985325) has multiple novel sequence insertions in the S protein compared to SARS-CoV (Urbani strain, GenBank accession no. AY278741), we aligned the S protein sequences of both viruses using a constraint-based alignment tool for multiple protein sequences (COBALT) [25]. We did not use MER-CoV for comparison because there is only 40% similarity between SARS-CoV-2 and MERS-CoV [26]. Similar to a previous report [21], we found sequence insertions (IS) in the SARS-CoV-2 S protein at four independent positions: IS1 “GTNGKTR”, IS2 “YYHK”, IS3 “HRSY”, and IS4 “NSPR” (Fig. 1A & B). To determine whether any of these sequence insertions constituted or resembled any protein motifs, such as an NLS, we analyzed the SARS-CoV-2 S protein in silico with the PSORT II web portal for NLS prediction [27]. We found that the SARS-CoV-2 glycoprotein contained an NLS of the “pat7” motifs, one of the three NLS motifs described above (Fig. S1). To our surprise, the NLS motif “PRRARSV” was present at the proposed polybasic site and was due to the fourth sequence insertion “NSPR” (Figs. 1A & B and S1). A widely reported furin consensus cleavage site motif is the canonical four amino acid motif R-X-[K/R]-R, although R-X-X-R is the minimal cleavage site on the substrate for successful furin cleavage [28, 29]. Due to the specificity of the amino acid motif, a furin cleavage motif is not expected to fulfill the characteristics of an NLS motif. However, the described furin cleavage site is constitutively within the NLS motif. Thus, whether furin cleavage destroys the function of the NLS motif is important to determine. As expected, the NLS in the S protein was unique to SARS-CoV-2 among human pathogenic beta-coronaviruses, as neither the SARS-CoV S protein nor the MERS-CoV S protein has an NLS (Fig. S1).

**Fig 1.**
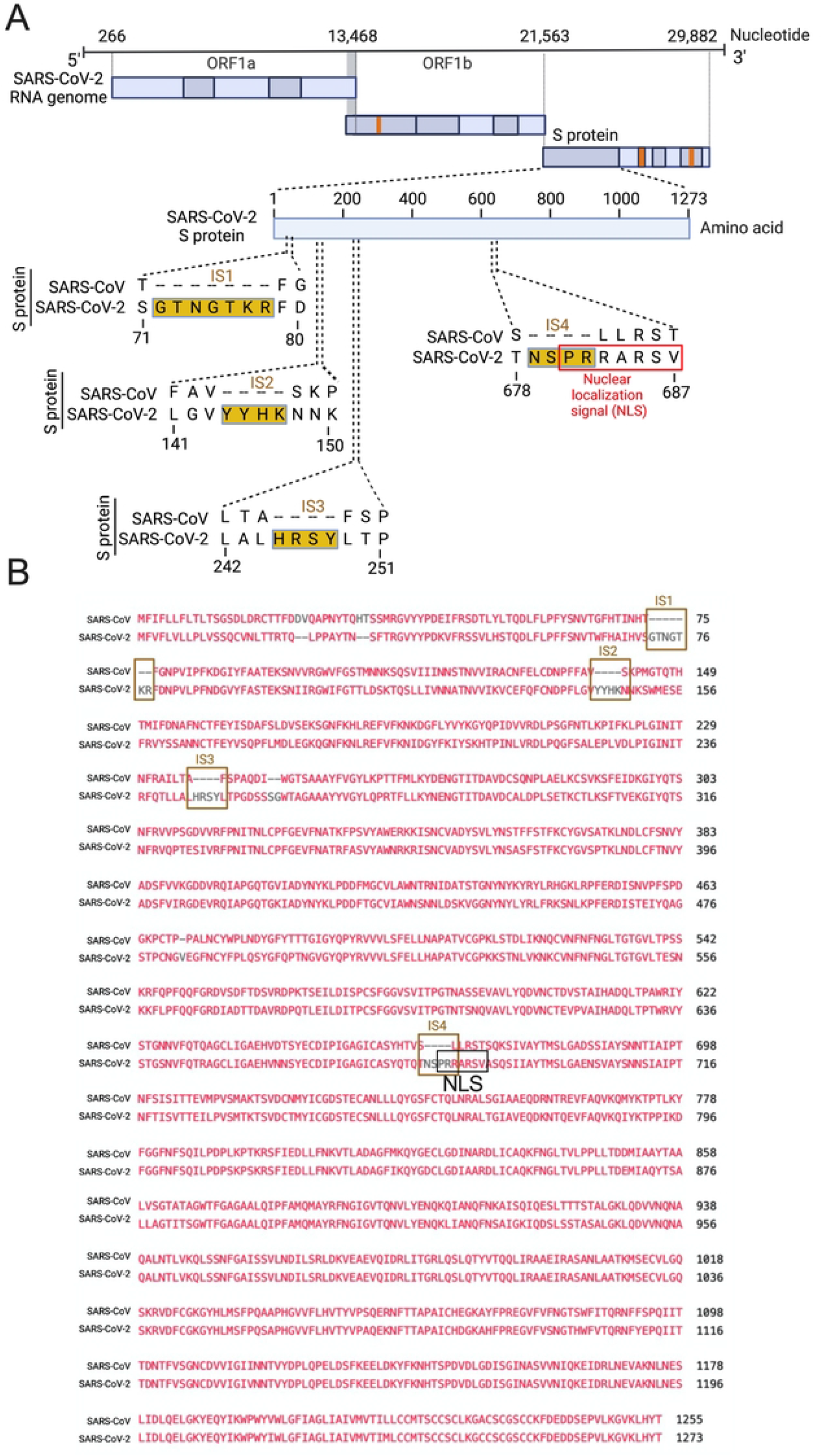
Only the SARS-CoV-2 S protein had an NLS motif “PRRARSV” due to a novel sequence insertion. **A**. Full-length SARS-CoV-2 genome (nucleotide) (USA/WA-CDC-WA1/2020 isolate, GenBank accession no. MN985325) and open reading frames (ORF) are shown at the top. The SARS-CoV-2 S protein amino acid sequence was aligned with SARS-CoV (Urbani strain, GenBank accession no. AY278741) by NCBI’s constraint-based multiple alignment tool COBALT [25], and the relative positions of four novel sequence insertions (ISs) are shown in the S protein ORF as follows: IS1: “GTNGKTR”, IS2: “YYHK”, IS3: “HRSY”, and IS4: “NSPR”. The fourth IS (NSPR) created a pat7 NLS “PRRARSV” in the S protein (shown in the red rectangle). **B**. The S protein ORF sequences between SARS-CoV-2 and SARS-CoV were aligned, and the Lined rectangles highlight the four novel insertions: IS1, IS2, IS3, and IS4. The IS4 “NSPR” created a pat7 NLS “PRRARSV” in the S protein (shown in the black rectangle).

### NLS-driven nuclear translocation of S protein (including S mRNA) occurs only in the SARS-CoV-2-infected airway epithelium

Although viral glycoprotein nuclear translocation is rare, NLS-driven protein nuclear translocation has already been established in different viral infections [30, 31]. Thus, it is important to determine whether the SARS-CoV-2 S protein translocates into the nucleus in addition to its canonical cell surface localization through the ER-Golgi pathway. We hypothesized that the S protein could translocate into the nucleus in SARS-CoV-2-infected cells via the identified NLS motif [17, 30, 31]. We infected highly differentiated pseudostratified airway epithelial cells (which mimics *in vivo* human airway epithelium) with SARS-CoV-2 at a multiplicity of infection (MOI) of 0.1 for four days. First, we confirmed the presence of S mRNA and S protein in a 5 µm section of formalin-fixed paraffin-embedded SARS-CoV-2-infected cells by RNAscope and immunofluorescence analysis. Despite the rarity of viral mRNA (or even positive-strand RNA virus genome) to be nuclear [32, 33], a recent study showed that SARS-CoV-2 mRNA accumulates in the nucleus of infected cells [34]. Our results showed that SARS-CoV-2 S mRNA was nuclear (Fig. 2 left panel and merged images in the right panel). To confirm the physical apposition between S mRNA and the nucleus by comparing their distributions in fluorescent images, we used the spot-to-spot colocalization function in Imaris image analysis software (Oxford Instruments).

**Fig 2.**
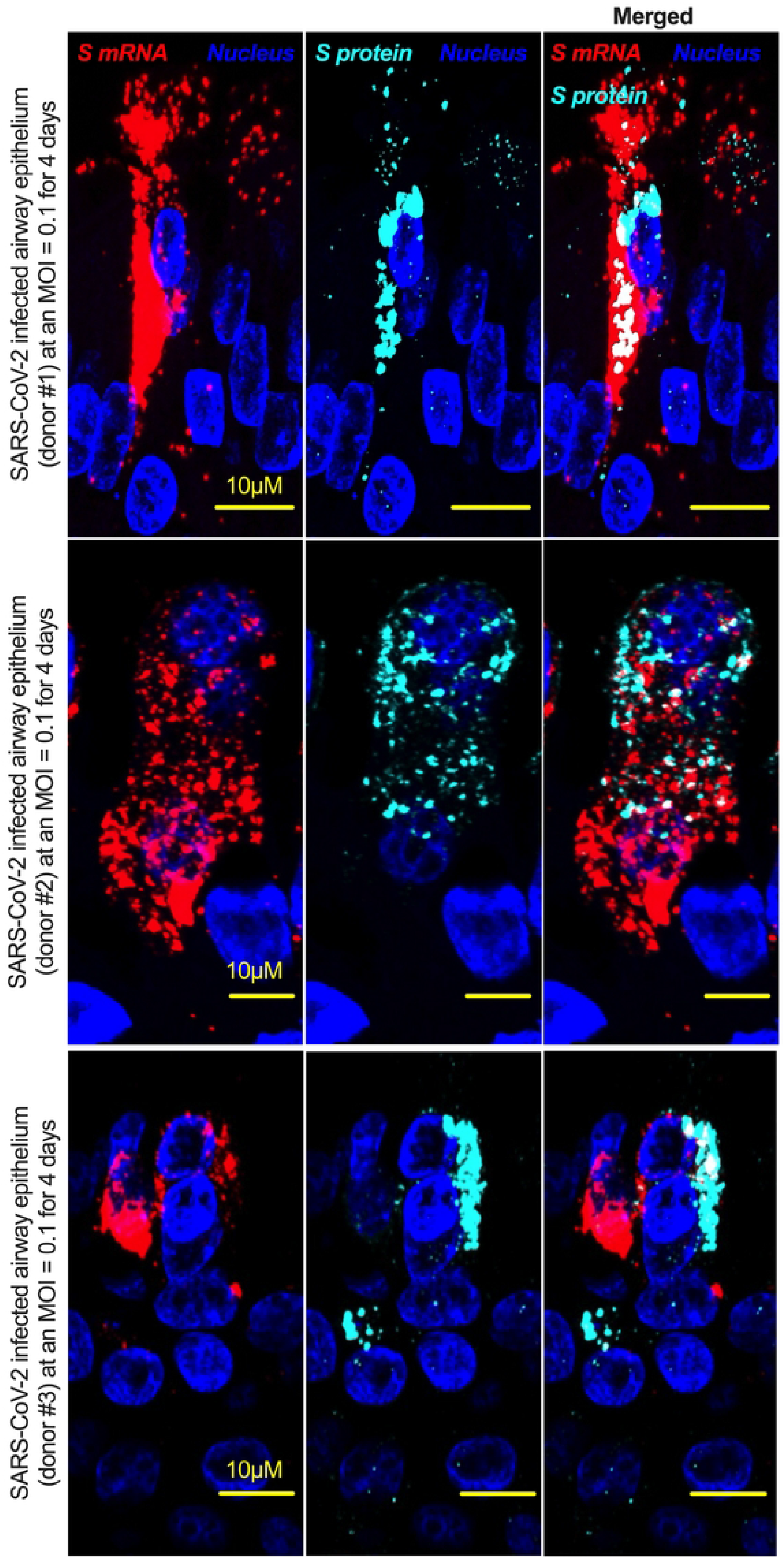
The intracellular distribution of S mRNA and S protein suggests nuclear translocation. Four-week **highly differentiated** pseudostratified airway epithelium was infected with SARS-CoV-2 at a MOI of 0.1 for four days, paraformaldehyde-fixed, paraffin-embedded, and sectioned at a thickness of 5 µm for immunohistochemistry (IHC) and slide preparation [43, 50]. A combined protocol of RNAscope and IHC was used to simultaneously detect S mRNA and S protein in the SARS-CoV-2-infected airway epithelium. S mRNA (red) was detected using a SARS-CoV-2 S mRNA probe for RNAscope, and S protein (cyan) was detected by an S protein-specific rabbit polyclonal antibody and a corresponding anti-rabbit secondary antibody for immunofluorescence (IFA) analysis. The nucleus (blue) was detected by DAPI staining. The images were taken under an Olympus confocal microscope using a 60x oil objective. The images represent multiple independent technical replicates from two independent experiments with different donors (experiment 1: donors 2 and 3 and experiment 2: donor 1). The scale bar is 10 µm.

We found that S mRNA was nuclear and abundant in the cytoplasm (Fig. S2, top panel, three donors). To avoid image artifacts, we imaged multiple independent slides of SARS-CoV-2-infected airway epithelium (from three independent donors) using at least two different high-end confocal microscopes. Additionally, we used at least two different image processing strategies to determine nuclear localization. Based on high-resolution imaging, we determined the subcellular distribution of S mRNA at the single-molecule and single-cell levels (Figs. 2 & 3A, S2 & S3). Importantly, we were able to determine S mRNA nuclear translocation not only inside the nucleus but also on the nuclear surface (Figs. 3A, 3C, and S3). The determination of S mRNA distribution and abundance showed that S mRNA subcellular localization spans from the inside and outer surface of the nuclear membrane to everywhere in the cytoplasm. We found that almost 90% of S mRNA was distributed in the cytoplasm, which was expected, as SARS-CoV-2 transcription and replication occur in the cytoplasm (Figs. 3 A & C, S3). Interestingly, less than 10% of S mRNA was detected at the nuclear surface, which could explain the transitionary stage of S mRNA before it enters the nucleus or the novel transnuclear-membrane translocation of S mRNA, which was examined and described later (Figs. 3 A & C, S3). In approximately 1% of instances, S mRNA successfully translocated into the nucleus (Figs. 3 A & C, S3). The nuclear translocation of S mRNA is highly unusual because there have been few previous reports of S mRNA nuclear translocation and no information on the mechanism of nuclear translocation. However, we have explored how S mRNA could translocate into the nucleus.

**Fig 3.**
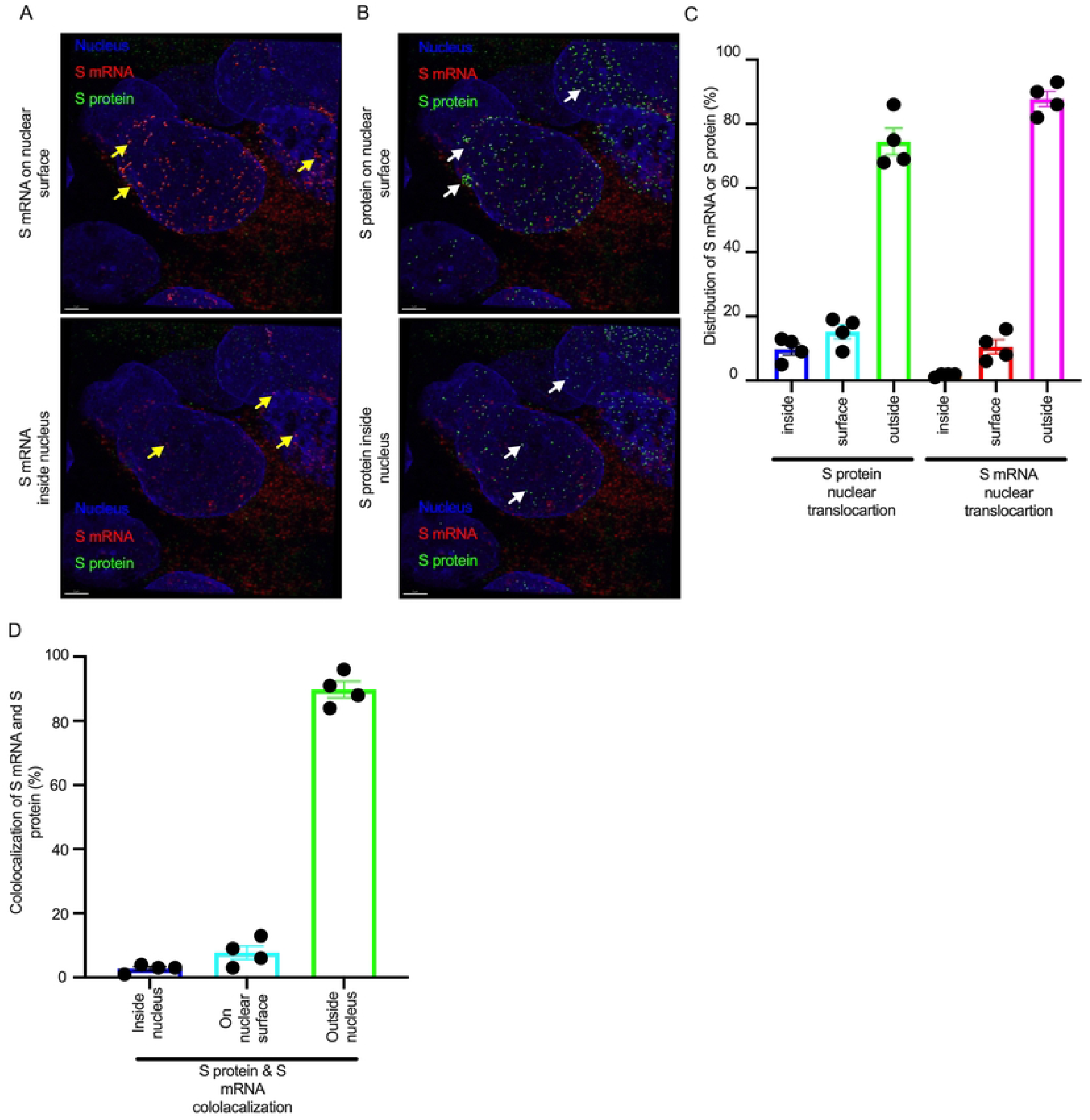
The nuclear translocation of S protein and S mRNA includes both the outer surface and inside of the nucleus. Separate slides (see Fig. 2) were imaged under a Leica Stellaris confocal microscope (Leica) using a 63x oil objective. The images were then deconvolved using Huygen Essential deconvolution software (Scientific Volume Imaging). Using the surface rendering function of an image processing IMARIS software. **A**. S mRNA (red) on the nuclear surface (top) and inside the nucleus (bottom). White arrows indicate S protein on the nuclear surface (top image) or inside the nucleus (bottom image). **B**. S protein (green) on the nuclear surface (top image) and inside the nucleus (bottom image). White arrows indicate S protein on the nuclear surface (top image) or inside the nucleus (bottom image). **C**. The total distribution of S mRNA and S protein in the cells. The data were obtained by combining multiple images from an independent experiment. **D**. The total colocalization between S mRNA and S protein in the cells. The data were obtained by combining multiple images from an independent experiment.

We investigated whether the S protein translocated into the nucleus in the SARS-CoV-2-infected airway epithelium. Consistent with the S mRNA data, we found that the S protein translocated into the nucleus and was abundant on the cellular surface through the cytoplasmic ER-Golgi pathway (Fig. 2, middle panel and merged images in the right panel). Based on high-resolution imaging, we determined that S protein nuclear translocation included both the inside of the nucleus and the nuclear surface (Figs. 3B, 3C, and S3). Similar to S mRNA quantification, we were able to quantify the distribution and abundance of S protein in the infected airway epithelium. We found that the S protein was distributed inside and in the outer membrane of the nucleus, the cytoplasmic ER-Golgi and the cell surface. We did not determine what portion of the S protein was localized inside and outside the cell surface. However, we quantified the subcellular distribution of the S protein inside the nucleus, outside the surface of the nucleus, and in the cytoplasm, which included cell surface expression because the S protein is a type 1 transmembrane glycoprotein. We found that approximately 75% of the S protein was distributed in sites other than the nucleus, including the cell surface and the cytoplasm, which was expected, as S protein translation and protein processing occur in the cytoplasm and via cytoplasmic ER-Golgi pathway, respectively (Figs. 3 B & C, S3). Interestingly, approximately 15% of the S protein was detected at the nuclear surface, which could explain the S protein transitionary stage before entering the nucleus or a novel transnuclear-membrane translocation of S protein, which was examined and described later (Figs. 3 A & C, S3). Interestingly, we found that a higher percentage of total S protein translocated into the nucleus than S mRNA (Figs. 3 A-C, S3). Although viral type-1 transmembrane glycoprotein translocation into the nucleus is rare, the NLS in the S protein is responsible for nuclear translocation. It was apparent that NLS-driven S protein nuclear translocation was SARS-CoV-2 specific, and a side-by-side infection experiment with both viruses showed that the S protein of SARS-CoV did not translocate into the nucleus (Fig. S4). As both S mRNA and S protein translocated into the nucleus, it is important to determine whether S mRNA and S protein colocalize in different subcellular sites.

### Colocalization of S mRNA and S protein in different subcellular locations in the SARS-CoV-2-infected airway epithelium

While we can explain S protein nuclear translocation due to the presence of an NLS motif in the amino acid sequence, we can only hypothesize that S mRNA nuclear translocation is possible due to a direct interaction between S protein and S mRNA, which can be explained by the colocalization between them. The SARS-CoV-2 N protein is an abundant RNA binding protein that is essential for viral genome packaging [35]. While the structural basis of N protein binding to single- or double-stranded RNA is known [36], there is no information about whether S protein binds to S mRNA. As we found similar intracellular distribution of both S mRNA and S protein, we hypothesized that S protein interacts with S mRNA to translocate the protein−mRNA complex to different subcellular locations, including the cytoplasm and nucleus, but not the cell surface. By examining the colocalization between the S protein and S mRNA, we could confirm the presence of the protein−mRNA complex in the SARS-CoV-2-infected airway epithelium. Here, we refer to colocalization as an association between S mRNA and S protein at different intracellular locations. Technically, two separate fluorescence molecules that emit different wavelengths of light are superimposed within an indeterminate microscope resolution. To determine whether S mRNA and S protein colocalize, we used a high-resolution imaging strategy. We quantified the colocalization on a percentage scale. We found that approximately 85% of the colocalization, which was the highest, was observed outside the nucleus (Fig. 3D). These data are consistent with the previously described spatial expression data of both S protein and S mRNA (Fig. 3A-C). As expected, lower and the lowest percentages of colocalization between the S protein and S mRNA were observed on the nuclear surface and inside the nucleus, respectively (Fig. 3D). We were able to pinpoint the colocalization site by using high magnification confocal imaging followed by image processing. Representatives of S mRNA and S protein colocalization in the cytoplasm (Fig 4, top two panels), on the surface of the nuclear membrane (Fig 4, middle two panels), or inside the nucleus (Fig 4, bottom two panels) are shown. We observed S protein and S mRNA colocalization in three subcellular locations, which was confirmed at the single-cell level in SARS-CoV-2-infected cells (Fig. S5). We also observed that S mRNA inside and on the nuclear surface was associated with the S protein, in contrast to the cytoplasmic S mRNA distribution with or without colocalization with the S protein (Figs. 4 and S5). Thus, S mRNA translocates into the nucleus through the S protein-S mRNA complex and is driven by the S protein (Figs. 4 and S5).

**Fig 4.**
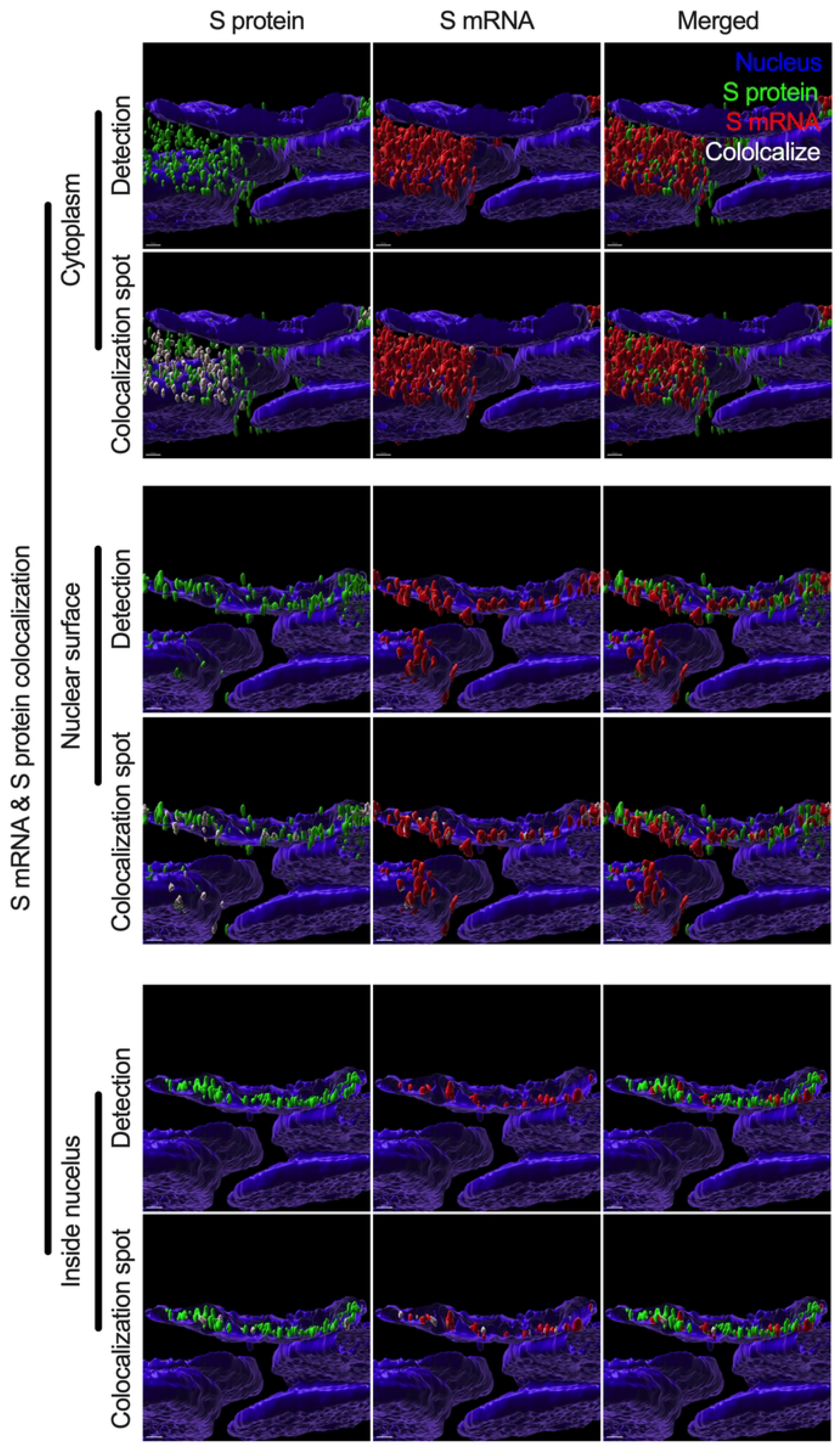
Colocalization between S mRNA and S protein inside infected cells. The images (see Fig. 3) were analyzed by using the surface rendering and colocalization features of IMARIS. S protein and S mRNA distribution and colocalization in the cytoplasm (top panel), on the nuclear surface (middle panel) and inside the nucleus (bottom panel). The specific region of colocalization is indicated by a white spot. Scale bar 0.5 µm.

### NLS-driven N protein nuclear translocation is common in SARS-CoV, MERS-CoV, and SARS-CoV-2 infections

There is no comprehensive information on SARS-CoV-2 N protein NLS motifs, and we already know that other pathogenic coronaviruses, particularly SARS-CoV [17, 37] and MERS-CoV [38] N proteins, have NLSs. Therefore, we searched for NLSs in the SARS-CoV-2 N protein by using the PSORT II. We found that the SARS-CoV-2 N protein has 7 NLSs covering all three types of NLS motifs (pat4: 2; pat7: 3, and bipartite: 2); however, the SARS-CoV N protein has 8 NLS motifs (pat4: 2; pat7: 4, and bipartite: 2) (Fig. S6). N protein translocation into the nucleus or at least the perinuclear region was confirmed (Figs. 5A and S7). However, SARS-CoV-2 N protein nuclear translocation was not as robust as SARS-CoV nuclear translocation (Figs. 5B and S7). The reduced nuclear translocation of the SARS-CoV-2 N protein is probably due to the absence of one pat7 NLS motif in the SARS-CoV-2 N protein compared to that in SARS-CoV (Figs. S7 and S8).

**Fig 5.**
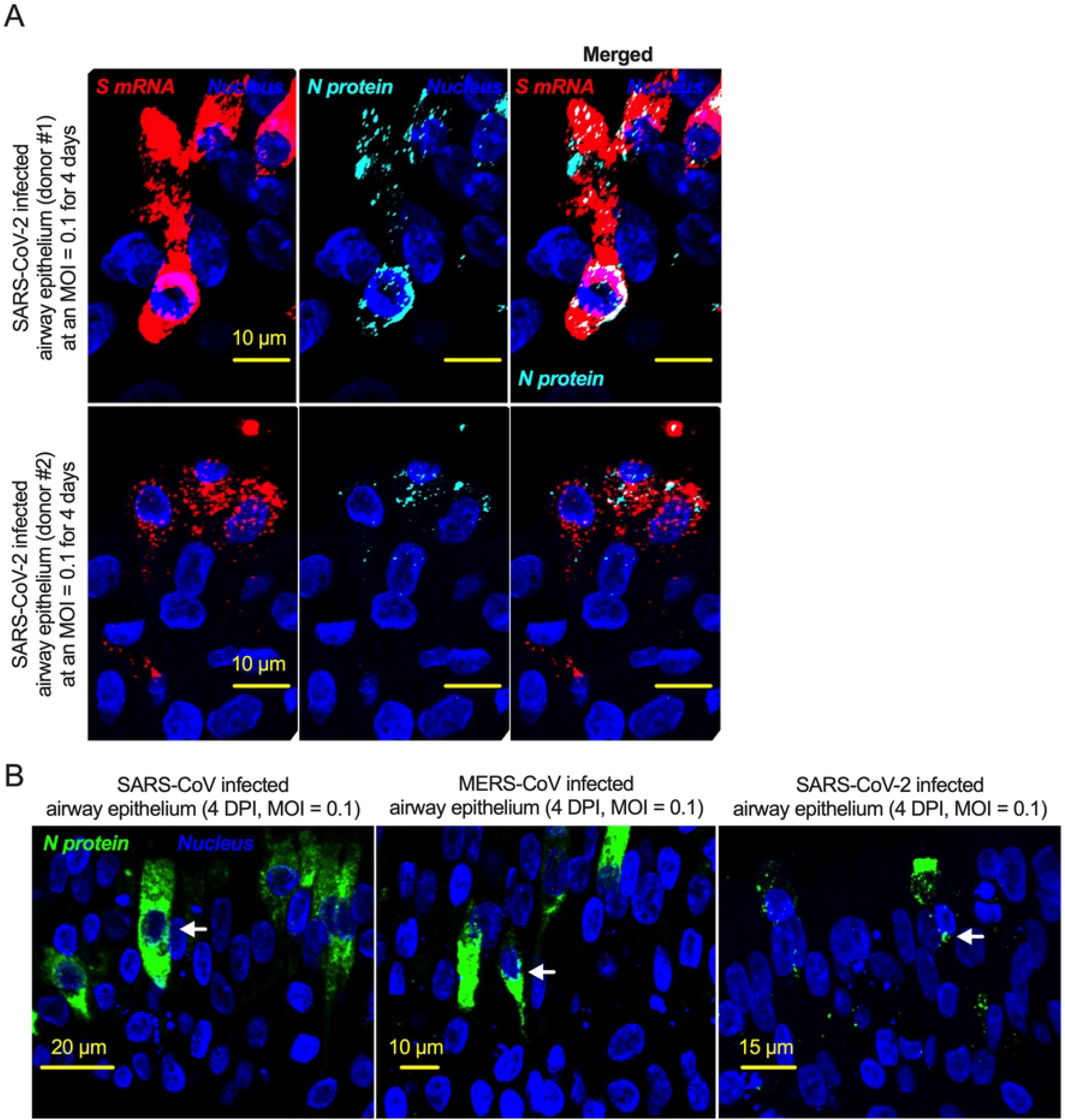
The nucleoproteins of SARS-CoV, MERS-CoV, and SARS-CoV-2 translocate into the nucleus. **A**. Four-week pseudostratified airway epithelium was infected with SARS-CoV-2 at a MOI of 0.1 for four days, fixed, paraffin-embedded, and sectioned at a thickness of 5 µm for immunohistochemistry slide preparation. Simultaneous detection of S mRNA (shown in red) and N protein (shown in cyan) on the same slide was performed by a combined detection protocol in RNAscope-based mRNA and immunofluorescence-based protein detection. An S mRNA-specific probe was used for RNAscope, and an N protein-specific rabbit polyclonal antibody and the corresponding anti-rabbit secondary antibody were used. The nucleus (shown in blue) was detected by DAPI staining. The images were taken under an Olympus confocal microscope using a 60x oil objective. The images represent multiple independent technical replicates from two independent, healthy donors (top row: donor #1 and bottom row: donor #2). The scale bar is 10 µm. **B**. Four-week pseudostratified airway epithelium was infected with SARS-CoV-2, SARS-CoV, or MERS-CoV at an MOI of 0.1 for four days. SARS-CoV-2 or SARS-CoV N protein was detected by an N protein-specific rabbit polyclonal antibody and corresponding anti-rabbit secondary antibody. Similarly, the MERS N protein was detected by MERS N protein-specific primary and corresponding secondary antibodies. The nucleus (shown in blue) was detected by DAPI staining. The images represent multiple independent technical replicates from one experiment (donor #1).

### SARS-CoV-2 S and N proteins’ interactions with genomic RNA

Based on a machine learning model, the SARS-CoV-2 RNA genome and sub-genomic RNAs can be translocated in the host cells’ mitochondrial matrix and nucleus [39]. Our results suggest that around 1% S mRNA translocated into the nucleus. S mRNA’s (potentially SARS-CoV-2 mRNA genome) subcellular localization maqy play a significant role in SARS-CoV-2 pathogenesis. To determine whether the SARS-CoV-2 genome interacts with either S protein or N protein, we in silico analyzed RNA-protein interactions using the RPISeq web portal, which offers the only sequence-based prediction model [40]. We found that both S and N protein binding probability to the SARS-CoV-2 genome scored exactly 1 (Dataset S1 and S2). SARS-CoV-2 N protein is an abundant RNA binding protein essential for viral genome packaging [35]. While the structural basis of N binding to single or double-stranded RNA is known [36], we found that S mRNA nuclear translocation aids via S protein. However, the mechanism of S protein binding to the mRNA or possibly positive-strand RNA genome is yet to be determined.

## Discussion

In the context of SARS-CoV, one of the controversies regarding the natural origin of SARS-CoV-2 is that its S gene has multiple novel sequence insertions. Zhang C. et al. analyzed the report by Pradhan et al. (withdrawn) [41] on the presence of four unique novel sequences in the SARS-CoV-2 S gene and showed that these four sequence insertions were not related to the receptor-binding domain (RBD) [21]. A recent study identified S gene novel sequence insertions among several key genomic features that differentiate SARS-CoV-2 from other beta-coronaviruses, particularly SARS-CoV and MERS-CoV [14]. The source and characterization of these sequence insertions have yet to be determined; however, the closest BLAST hit of these sequences is bat coronavirus RaTG13 [42]. Similar to a previous report [21], we found four multiple sequence insertions in the SARS-CoV-2 S protein: IS1 “GTNGKTR”, IS2 “YYHK”, “HRSY”, and IS4 “NSPR” (Fig. 1).

Here, we showed that the fourth novel sequence insertion in the S gene was an NLS and resulted in nuclear translocation of the S protein, which not only complemented previous in silico findings [14] but also identified a novel pathogenic genomic feature of the S gene. Interestingly, the fourth significant insertion has received attention due to the description of a polybasic site “RRAR”, which may contribute to increased serin protease-driven entry of SARS-CoV-2 [19] and is implicated in broader tropism and/or enhanced viral transmissibility compared to SARS-CoV [20]. However, we found that the IS4 “NSPR” created a pat7 NLS “PRRARSV” in the S protein, which was unique to SARS-CoV-2. We first reported that the S protein translocated into the nucleus in the SARS-CoV-2-infected airway epithelium, which is an appropriate lung model for studying respiratory virus infection in vitro [2, 43]. Our results confirmed that the SARS-CoV-2 S protein was a unique addition to the list of viral proteins that possess NLSs and consequently translocate into the nucleus of infected cells [17, 30, 31, 44]. Among coronaviruses, SARS-CoV-2 S protein is the first type-1 transmembrane glycoprotein that translocates into the nucleus. Vesicular stomatitis virus (VSV), which is a negative sense RNA virus, has a glycoprotein that translocates to the nucleus as well. A study by the University of Illinois at Urbana Champaign showed exactly how the glycoprotein on VSV was able to travel to the nucleus of hamster kidney cells [45].

NLS-driven S protein nuclear translocation is a novel pathogenic feature of SARS-CoV-2 infection compared to other pathogenic coronaviruses. However, the pathogenic contribution of the S protein’s NLS motif to virus-induced pathophysiology is yet to be determined. Our results suggested that the S protein translocated into the nucleus due to the NLS, which also raised two important points. First, we investigated whether the proposed polybasic site “RRAR” could itself be an NLS motif. The answer was that the proposed polybasic site was not an NLS motif because an NLS is a well characterized and predefined amino acid sequence motif [15-17]. Additionally, the amino acid sequence of the probable sequence insertion “NSPR” [21] was also not an NLS but was part of the P7 “PRRARSV” NLS. Thus, the inserted sequence creates the NLS in the S protein of SARS-CoV-2 and may make SARS-CoV-2 unique among human pathogenic coronaviruses.

The second important point was whether the NLS motif was functional in the context of the described polybasic site at the S1/S2 boundary. All type-1 transmembrane glycoproteins are processed through the ER-Golgi pathway before signal peptide-driven cellular surface localization. The proposed polybasic site was functional (availability to proteases) when the S protein was on the virion for host cell entry. A fully posttranslationally processed S protein surface translocation could also provide a polybasic site to be processed by furin cleavage. However, there is no information on the availability or usability of the S protein’s polybasic site by furin proteases in the cytoplasm before virus assembly. Thus, the NLS is functional in SARS-CoV-2-infected cells, and the polybasic site only functions during the viral entry step. The NLS is obviously functional in infected cells, and no furin cleavage at the polybasic site is necessary other than for viral entry. Our results confirmed that the S protein NLS motif was functional in SARS-CoV-2-infected cells. Although mutating the polybasic site (which also mutated the NLS) may impact viral S protein function in vitro, the result will not confirm or deny that one is more important than the other between the polybasic site and the NLS. While our result does provide direct evidence for the presence of the NLS motif and nuclear translocation of the S protein, our results do not confirm nor deny that the NSPR sequence has a natural origin. Instead, our results showed that the inserted sequence NSPR was a functional NLS motif, which increased the intracellular distribution of the S protein, including novel nuclear translocation. The novel nuclear translocation of the SARS-CoV-2 S protein suggests that: 1. the nuclear translocation of the S protein reduces its surface expression, but whether it contributes to evading host immune recognition remains to be determined; and 2. the colocalization of the S protein with S mRNA suggests that the S protein has an RNA binding motif, which remains to be determined. One of the important ways of confirming a functional NLS motif is to use site-directed mutational analysis. Plasmid-driven transient expression of S protein in the human lung airway A549 cell line and primary normal human branchial epithelial cells showed robust S expression but was toxic to the cells. Therefore, the success of site-directed mutational analysis of the S protein in a transient expression system is doubtful and the characterization of NLS by a mutational analysis is yet to be determined. Thus, our novel findings emphasize further research on the NLS motif of the SARS-CoV-2 S protein.

One of the most important findings in our study was the simultaneous detection of the different spatial distributions of S protein and S mRNA at the single-molecule level in a single infected cell. We confirmed that S mRNA translocated into the nucleus by image analysis of the colocalization of S mRNA with nuclear staining. The SARS-CoV-2 N protein has already been shown to bind to RNA [46]. There was no information available confirming whether the S protein could bind to S mRNA for nuclear translocation. Our results revealed that S mRNA nuclear translocation was mediated by the S protein because S mRNA nuclear translocation was always associated with the S protein. For example, S mRNA colocalized with the S protein inside and outside the surface of the nucleus. Although the primer-probe was designed to target S mRNA, the SARS-CoV-2 positive-strand RNA genome (whole or partial) can be targeted by the same probe due to the sequence similarity between S mRNA and the whole or partial genome. Thus, our results lack sufficient detail contributing to the discussion of the controversial scientific topic of whether there is any possibility of SARS-CoV-2 genome integration into the host DNA [47, 48]. Additionally, one of the significant differences in the S protein sequences of SARS-CoV and SARS-CoV-2 is the pat7 NLS motif. Whether S protein expression by the current vaccine platforms causes suboptimal expression of S protein on the cell surface due to the NLS remains to be determined [49].

In conclusion, the SARS-CoV-2 S protein has a functional pat7 NLS “PRRARSV”, that results in one out of four S proteins translocating into the nucleus in infected cells. S Protein appears to shuttle S mRNA (possibly the genome) into the nucleus as well. Thus, the NLS of the S protein may contribute to the evasion of the host immune response and is a novel pathogenic feature of SARS-CoV-2.

## Materials and Methods

### Cells and viruses

Primary normal human bronchial epithelial (NHBE) cells from healthy adults and high-risk adults (deidentified) were obtained from Dr. Kristina Bailey at the University of Nebraska Medical Center (UNMC) (Omaha, NE) under an approved material transfer agreement (MTA) between the University of North Dakota (UND) and UNMC (Omaha, NE). The protocol for obtaining cells was reviewed by the UNMC IRB and was determined to not constitute human subject research (#318-09-NH). In this study, we used cells from five donors: nonsmoker healthy adults (donors #1 and #2) and adult with chronic obstructive pulmonary disease (COPD) (donor #3). The protocols for subculturing primary NHBE cells were published previously [2, 43, 50]. SARS-CoV-2 (USA/WA-CDC-WA1/2020 isolate, GenBank accession no. MN985325; kindly provided by CDC), SARS-CoV (Urbani strain, GenBank accession no. AY278741; kindly provided by Rocky Mountain Laboratories (RML), NIAID, NIH), and MERS-CoV (GenBank accession no. NC_019843.3; kindly provided by the Department of Viroscience, Erasmus Medical Center, Rotterdam, The Netherlands) were used for *in vitro* infections described below.

### In silico analysis

We have used open-source web portals for different *in silico* analyses. 1. Constraint-based alignment tool for multiple protein sequences (COBALT) (https://www.ncbi.nlm.nih.gov/tools/cobalt/re_cobalt.cgi) was used for multiple sequence alignment. 2. PSORT II (https://psort.hgc.jp/form2.html) was used for NLS prediction. 3. The RPIseq web portal (http://pridb.gdcb.iastate.edu/RPISeq/) was used for RNA−protein interactions.

### Highly differentiated pseudostratified bronchial airway epithelium

The protocols for differentiating primary NHBE cells to form a pseudostratified bronchial airway epithelium were published previously [2, 43, 50]. Briefly, Transwells (6.5 mm) with 0.4-µm-pore polyester membrane inserts (Corning Inc.) were coated with PureCol (Advanced BioMatrix) for 20 min before cell seeding. NHBE cells (5×10^4) suspended in 100 µl of complete airway epithelial cell (cAEC) medium [AEC medium (Promocell) + SupplementMix (Promocell) + 1% penicillin−streptomycin (V/V) (Thermo Fisher Scientific) + 0.5% amphotericin B (V/V) (Thermo Fisher Scientific)] were seeded in the upper chamber of the Transwell. Then, 500 µl of cAEC medium was added to the lower chamber of the Transwell. When the cells formed a confluent layer on the Transwell insert, the cAEC medium was removed from the upper chamber, and in the lower chamber, the cAEC medium was replaced with complete ALI medium [PneumaCult-ALI basal medium (Stemcell Technologies Inc.) + with the required supplements (Stemcell Technologies) + 2% penicillin−streptomycin (V/V) + 1% amphotericin B (V/V)]. The complete ALI medium in the lower chamber was changed every day. The upper chamber was washed with 1x Dulbecco’s phosphate-buffered saline (DPBS) (Thermo Fisher Scientific) once per week initially but more frequently when more mucus was observed during later days. All cells were differentiated for at least four weeks at 37°C in a 5% CO_2_ incubator. We observed motile cilia in the differentiated airway epithelium similar to previously described [50].

### Viral infection

All viral infection experiments were conducted in the high biocontainment facility at RML, NIAID, NIH, Hamilton, MT. After approximately 3 weeks, the differentiated airway epithelium on Transwells was shipped to RML in an optimized transportation medium [2, 50], and the recovered cells were maintained in complete ALI medium for approximately one week before infection. For infection, the airway epithelium on Transwells was washed with 200 µl of 1x PBS to remove mucus and were infected on the apical site with SARS-CoV-2, MERS-CoV, or SARS-CoV at a MOI of 0.1 in 100 µl 1x PBS for 1 hour (at 37°C with 5% CO_2_). For mock infection, the Transwells were similarly incubated with 100 µl 1x PBS without virus. The viral inoculum was then removed, and the epithelium on the Transwell was washed twice with 200 µl of 1x PBS. Complete ALI medium (1000 µl) was added to the lower chamber of each Transwell, and the upper chamber was kept empty. Mock-infected and virus-infected Transwells were incubated for 4 days at 37°C in an incubator with 5% CO_2_ [2].

### Paraformaldehyde (PFA) fixation and paraffin embedding

At 4 days postinfection (DPI), 200 μl of 1x PBS was added to the apical site of the Transwell for washing before PFA fixation. In the basal side of the Transwell inserts was 200 μl of 1x PBS. For PFA fixation, 200 μl of 4% PFA (Polysciences) was added to the upper chamber of the Transwells and incubated for 30 min, and the Transwells were further maintained overnight in 4% PFA prior to removal from high biocontainment. The PFA fixation protocol was approved as an inactivation method for coronaviruses by the RML Institutional Biosafety Committee. The PFA-fixed airway epithelium was paraffin-embedded and sectioned at a thickness of 5 µm for slide preparation as previously described [43].

### Simultaneous detection of S mRNA and S protein

Slides with 5 µm sections were first deparaffinized by incubation in a Coplin jar as follows: 1. Histo-Clear for 5 min, two times; 2. 100% ethanol for 5 min, three times; 3. 95% ethanol for 5 min, 4. 70% ethanol for 5 min, and 5. distilled water for 5 min. The deparaffinized slides were immediately incubated in 0.5% Triton X-100 in 1x PBS for 30 min. The slides were washed three times with 1X PBST (1x PBS with Tween 20) or 1x PBS for 5 min. A hydrophobic barrier was drawn around the 5 µm section on the slides by using an Immedge Hydrophobic Barrier Pen. To reduce nonspecific antibody binding, the section was blocked with 10% goat serum (Vector Laboratories) in 1x PBST for 2 hours at 4°C. The slides were then incubated with viral protein-specific primary antibody solution in 1x PBST (e.g., SARS-CoV/SARS-CoV-2 S protein-specific rabbit polyclonal antibody at a 1:100 dilution, SARS-CoV/SARS-CoV-2 N protein-specific mouse monoclonal antibody at a 1:100 dilution, or MERS-CoV N protein-specific mouse monoclonal antibody at a 1:100 dilution) overnight at 4°C. The slides were then incubated with the corresponding secondary antibody solution (anti-mouse or anti-rabbit AF488 or AF647, Thermo Fisher Scientific) in 1x PBST for 2 hours at room temperature. We then stained the nuclei with DAPI reagent (Advanced Cell Diagnostics) or used RNAscope multiplex V2 to detect SARS-CoV-2 S mRNA (Probe V-nCoV2019-S) according to the manufacturer’s instructions (Advanced Cell Diagnostics). The sections were mounted on Tech-Med microscope slides (Thomas Fisher Scientific) using ProLong-Gold antifade mounting medium (Thermo Fisher Scientific).

### Imaging and image analysis

The images were taken under an Olympus FluoView laser scanning confocal microscope (Olympus FV3000) enabled with a 60X objective (Olympus), a Leica Stellaris confocal microscope (Leica Microsystem) using a 63x oil objective or a Leica DMI8 epifluorescence microscope (Leica Microsystem). The images were then deconvolved using Huygen Essential deconvolution software (Scientific Volume Imaging). The surface rendering function of Imaris image processing software (Oxford Instruments) was used. The images were also analyzed for spot-to-spot colocalization by Imaris. Where applicable, images taken under a Leica DMI8 microscope were processed using 3D deconvolution and 3D view modules in LASX software (Leica Microsystem). For figure preparation, Prism version 9 (GraphPad) and Adobe Photoshop (Creative Cloud) software were used.

---

**Fig S1. NLS prediction in the S protein of pathogenic coronaviruses**. All categories of NLS motifs were searched in the S protein sequence using the web-based program PSORT II (https://psort.hgc.jp/form2.html) [27] for the S protein ORF amino acid sequence of SARS-CoV-2 (USA/WA-CDC-WA1/2020 isolate, GenBank accession no. MN985325) (Query 1), SARS-CoV (Urbani strain, GenBank accession no. AY278741) (Query 2), or MERS-CoV (GenBank accession no. NC_019843.3) (Query 3).

**Fig S2. Detection of the nuclear translocation of S mRNA** and **S protein**. Confocal images of SARS-CoV-2-infected airway epithelium (described in Fig 2) were analyzed for spot-to-spot colocalization using Imaris image analysis software (Oxford Instruments). The left panel shows the confocal images, the middle panel shows spot-to-spot colocalization, and the right panel shows merged confocal images and spot-to-spot colocalization. Spot-to-spot colocalization between the nucleus and S protein or S mRNA is indicated by a different color. The images represent multiple independent cross sections of the SARS-CoV-2-infected airway epithelium (from 3 independent donors).

**Fig S3. The translocation of S mRNA and S protein includes both the inside and outer surface of the nucleus**. From the images shown in Fig 3, the signals of S mRNA and S protein were plotted in the graph by Imaris image analysis software. The distance and intensity of all S mRNA or S protein from the nuclear surface (considered 0) were plotted. A negative value indicates that S mRNA or S protein resides inside the nucleus. The higher the negative value is, the farther the distance from the nuclear surface. In contrast, a positive value indicates that S mRNA or S protein resides on the nucleus surface and beyond in the cytoplasm. The higher the positive value is, the farther the distance from the nuclear surface.

**Fig S4. The SARS-CoV S protein does not translocate into the nucleus**. A four-week pseudostratified airway epithelium was infected with SARS-CoV at an MOI of 0.1 for four days, fixed, paraffin-embedded and sectioned at a thickness of 5 µm for immunohistochemistry slide preparation. S protein (shown in cyan) was detected by immunofluorescence-based protein detection using SARS-CoV/SARS-CoV-2 S specific rabbit polyclonal primary antibody and anti-rabbit secondary antibody. The confocal image was analyzed for spot-to-spot colocalization using Imaris image analysis software. The left panel shows a confocal image, the middle panel shows spot-to-spot colocalization, and the right panel shows merged confocal images and spot-to-spot colocalization. The images represent multiple independent cross sections of SARS-CoV-infected airway epithelium (at least two donors).

**Fig S5. S mRNA and S protein colocalization was spatially evident in all possible ways inside the infected cell**. The confocal images shown in Figs. 3 & 4 were further visualized at a higher magnification to detect S mRNA and S protein colocalization spatially. S protein and S mRNA distribution and colocalization in the cytoplasm (top panel), on the nuclear surface (middle panel) and inside the nucleus (bottom panel). The specific region of colocalization is indicated by a white spot. The colors were made translucent to show colocalization. Scale bar 0.2 µm.

**Fig S6. NLS motif prediction in the N protein of pathogenic coronaviruses**. All categories of NLS motifs were searched in the N protein sequence using the web-based program PSORT II (https://psort.hgc.jp/form2.html) [27] for the N protein ORF of SARS-CoV-2 (USA/WA-CDC-WA1/2020 isolate, GenBank accession no. MN985325) (Query 1), SARS-CoV (Urbani strain, GenBank accession no. AY278741) (Query 2), or MERS-CoV (GenBank accession no. NC_019843.3) (Query 3).

**Fig S7. Nuclear translocation of the N protein of pathogenic coronaviruses**. Four-week pseudostratified airway epithelium was infected with SARS-CoV-2, SARS-CoV, or MERS-CoV at an MOI of 0.1 for four days, fixed, paraffin-embedded and sectioned at a thickness of 5 µm for immunohistochemistry slide preparation. SARS-CoV-2 or SARS-CoV N protein (green) was detected by a SARS-CoV/SARS-CoV-2 N protein-specific antibody. Similarly, the MERS N protein (green) was detected by the MERS N protein-specific antibody. The nucleus (shown in blue) was detected by DAPI staining. The confocal images were analyzed for spot-to-spot colocalization. The left panel shows a confocal image, the middle panel shows spot-to-spot colocalization, and the right panel shows merged confocal images and spot-to-spot colocalization. Spot colocalization between the nucleus and N protein is indicated by a different color. The images represent multiple independent technical replicates from at least one independent experiment for one donor (donor #1).

**Fig S8. NLS motif distribution in the N protein in different pathogenic coronaviruses**. The sequences of the N protein of the SARS-CoV-2 N protein (nCoV-WA1-2020, GenBank accession no. MN985325), SARS-CoV N protein (Urbani Strain, GenBank accession no. AY278741), and MERS-CoV N protein (HCoV-EMC/2012, GenBank accession no. NC_019843) by NCBI’s constraint-based multiple alignment tool COBALT [25]. All categories of NLS motifs are shown in the colored rectangle box: pat4: green; pat7: blue; bipartite 1: black; bipartite 2: orange.

**S1 dataset 1**. Prediction of SARS-CoV-2 S protein and genome interaction

**S2 dataset 2**. Prediction of SARS-CoV-2 N protein and genome interaction

## Acknowledgments

We are thankful to the MARC U-STAR program at UND for supporting the undergraduate students Sarah Sattar and Kailey Jerome. We are grateful to Dr. Jaspreet K. Osan for helping with primary cell culture work. We are also grateful to Heinz Feldmann of the Laboratory of Virology, NIAID, NIH for support with material and infections. In addition, we thank the Microscopy Core (UND, Grand Forks), which is funded by NIH P20GM103442, of the INBRE program for providing access to an Olympus FV300 confocal microscope. Histological services were provided by the UND Histology Core, which is supported by the NIH/NIGMS awards P20GM113123, U54G M128729, and UND SMHS funds. We also thank the Imaging Core (UND, Grand Forks), which is funded by NIH P20GM113123, NIH U54GM128729, and UNDSMHS funds, for IMARIS image analyses. This work was funded by the NIH/NIGMS awards P20GM113123 and T34GM122835, VA grant 101-BX005413 and partially by the Intramural Research Program, NIAID, NIH. The content is solely the responsibility of the authors and does not necessarily represent the official views of the NIH.

## Author contributions

M.M. conceived the project and designed all the experiments. K.B. provided the primary cells. F.F. performed the viral infection work. K.J., S.S. and M.M. performed all staining for detection. M.M. and S.S. generated the microscopic images. J.K. and M.M. processed and quantified images. M.M. analyzed (in silico) the viral genome and protein sequences. M.M. prepared the figures and wrote and edited the manuscript.

